# Increased microtubule lattice spacing correlates with selective binding of kinesin-1 in cells

**DOI:** 10.1101/2022.05.25.493428

**Authors:** Leanne de Jager, Klara I. Jansen, Lukas C. Kapitein, Friedrich Förster, Stuart C. Howes

## Abstract

Within the cell cargo is transported via motor proteins walking along microtubules. The affinity of motor proteins for microtubules is controlled by various layers of regulation like tubulin isoforms, post-translational modifications and microtubule associated proteins. Recently, the conformation of the microtubule lattice has also emerged as a potential regulatory factor, but to what extent it acts as an additional layer of regulation has remained unclear. In this study, we used cryo-correlative light and electron microscopy to study microtubule lattices inside cells. We find that, while most microtubules have a compacted lattice (∼41 Å), a significant proportion of the microtubule cores have expanded lattice spacings and that these lattice spacings could be modulated by the microtubule stabilizing drug Taxol. Furthermore, kinesin-1 predominantly binds microtubules with a more expanded lattice spacing (∼41.6 Å). The different lattice spacings present in the cell can thus act as an additional factor that modulates the binding of motor proteins to specific microtubule subsets.

## Introduction

Microtubules are highly dynamic, polarized cytoskeletal structures along which motor proteins move to transport cargos throughout the cell. To achieve efficient cargo transport, the binding of motor proteins to specific subsets of microtubules is highly regulated. The multiple layers of regulation and modifications that mark the different populations of microtubules and influence the downstream behaviour of motor proteins have together been termed the tubulin code (Verhey and Gaertig, 2007). Many of the components of this code, like tubulin isoforms, post-translational modifications (PTMs) and microtubule associated proteins (MAPs), have been identified (Gadadhar et al., 2017; Janke and Magiera, 2020; Roll-Mecak, 2020). However, the tubulin code is potentially still incomplete as it does not explain all motor protein behaviour observed *in vivo*.

Specifically, the motor protein kinesin-1 is known to be regulated by various components of the tubulin code (Janke and Magiera, 2020). For example, the binding properties of kinesin-1 are modulated by MAPs. Binding of MAP7 leads to activation of kinesin-1, namely by increasing its run length, while tau has the opposite effect (Ferro et al., 2022; Hooikaas et al., 2019; Monroy et al., 2018). Furthermore, kinesin-1 binds with greater affinity to a subset of stable microtubules (typically enriched in the PTMs acetylation and detyrosination) than to microtubules without these modifications (Cai et al., 2009; Dunn et al., 2008; Guardia et al., 2016; Konishi and Setou, 2009; Liao and Gundersen, 1998; Tas et al., 2017). However, *in vitro* experiments to date, where different levels of these PTMs were investigated, have not reproduced the behaviour of kinesin-1 that is observed *in vivo* (Kaul et al., 2014; Sirajuddin et al., 2014; Walter et al., 2012). In addition, upon treatments that lead to acetylation or detyrosination of most cellular microtubules, kinesin-1 still binds to a subset of stable microtubules (Jansen et al., 2021). This indicates that the tubulin code for kinesin-1 is not completely understood.

An additional layer of regulation might come from the structure of the microtubule itself, as several studies have shown that proteins are sensitive to its lattice spacing and/or nucleotide state (Manka and Moores, 2018a; Zhang et al., 2018). Upon polymerization of free αβ-tubulin dimers into microtubules, GTP bound to β-tubulin is hydrolysed to GDP and the structural conformation of the αβ- tubulin dimer changes (Alushin et al., 2014). This GDP-bound state, which makes up the bulk of the microtubule core, generally has a compacted average monomer spacing of ∼41 Å that is distinct from that of the GTP-bound state at the tip (GTP-cap), which has an expanded lattice spacing of ∼42 Å as determined for hydrolysis deficient mutants and the slowly-hydrolysable GTP analogue guanylyl-(α,β)- methylene-diphosphonate (GMPCPP) (Hyman et al., 1995; LaFrance et al., 2021). Besides the nucleotide state, drugs, like the anti-cancer drug Taxol, can alter the lattice spacing of the microtubule (Kellogg et al., 2017). Some MAPs can differentiate between these different lattice spacings. For example, end binding proteins and doublecortin preferentially bind to the GTP-cap of the microtubule (Bechstedt and Brouhard, 2012; Tirnauer and Bierer, 2000), while Tau (Duan et al., 2017) and MAP2 preferentially bind to microtubules in the GDP-bound state (Siahaan et al., 2021). A similar lattice sensitivity might partially explain the observed microtubule-subset specificity of kinesin-1.

*In vitro* data show that kinesin-1 binds expanded microtubules with a higher affinity than microtubules in the compacted state and that kinesin-1 binds microtubules assembled with GMPCPP at a higher frequency than GDP-bound microtubules (Shima et al., 2018). Moreover, GDP-microtubules, when heavily decorated with kinesin-1, were reported to be more expanded than undecorated microtubules (Shima et al., 2018) and saturating concentrations of kinesin-1 induced microtubule lattice expansion (Peet et al., 2018). However, these experiments were all performed in reconstituted systems using brain tubulin that contains a diverse mixture of PTMs and tubulin isoforms (Schwarz et al., 1998). Additionally, the observed effects depended on the kinesin-1 concentrations used, but the relationship to *in vivo* concentrations was not established. It is thus still unclear to what extent the microtubule lattice spacing plays a regulatory role within the cell.

To investigate if subsets of microtubules with varying lattice states exist within cells, and to determine if kinesin-1 preferentially binds a particular lattice, we established a cryo-correlative light and electron microscopy (cryo-CLEM) workflow. We show here that microtubules can have diverse lattice spacings in cells and that Taxol treatment greatly expands the lattice. Furthermore, we find that kinesin-1-decorated microtubules are predominately expanded in cells.

## Results and discussion

### Microtubule lattice spacings within cells are diverse but mainly compacted

To examine the lattice spacing of microtubules in cells, we cultured U2OS cells on EM grids, vitrified them, and prepared ∼150 nm thick lamellae using cryo-focused ion beam (FIB) milling for subsequent cryo-electron tomography imaging (Rigort et al., 2012). To measure the lattice spacing of cellular microtubules, we used a layer line approach based on Fourier analysis of microtubule segments (Figure 1A). Building from typical 2D analysis of *in vitro* microtubules (Mandelkow et al., 1977), the protocol masks out microtubule density from surrounding cellular material and aligns the segments in 3D (Figure 1B), prior to projection and computation of the 2D Fourier transforms (Figure 1C and Supplemental Figure 1). Summed power spectra of the aligned microtubule segments were then used to measure the lattice periodicity based on maxima in a line profile plot (Figure 1G).

**Figure 1.**
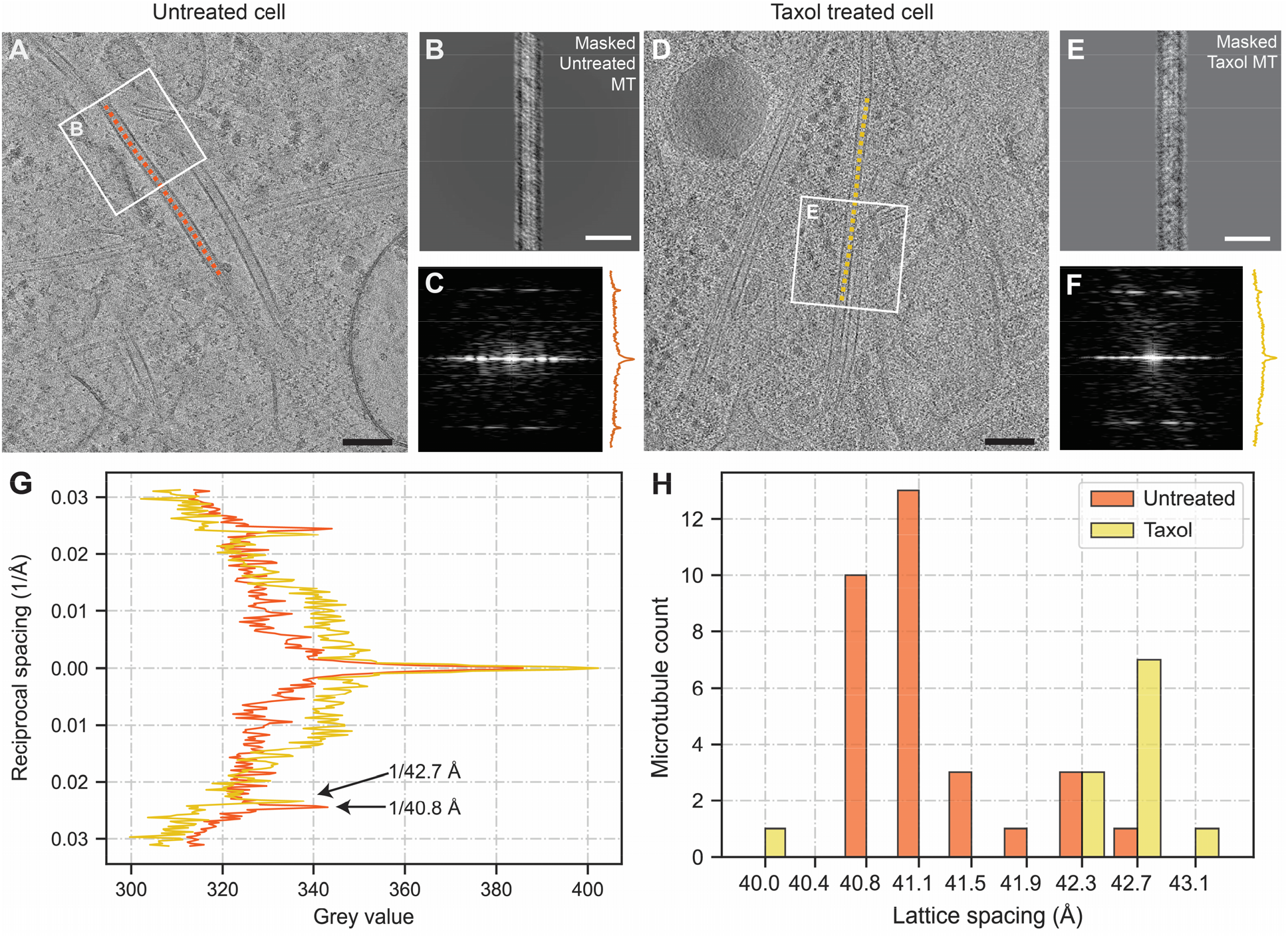
Microtubule lattice spacings within cells are diverse and expand upon Taxol treatment. (**A**) Tomogram slice (thickness 10 nm) showing the selected microtubule (MT) backbone of a MT in an untreated U2OS cell (orange). (**B**) Aligned and masked MT segment from the MT shown in A. (**C**) Power spectrum of the MT segment and its layer line plot from the MT shown in A. (**D-F**) Similar MT layer line analysis performed on Taxol-treated U2OS cells (yellow). (**G**) Layer line plot of the summed power spectra of segments from the same MT in A (orange, untreated) and D (yellow, Taxol), arrows indicate the location of the layer line peak and its related lattice spacing. (**H**) Histogram showing the untreated (orange, N=31, 12 tomograms) and Taxol-treated (yellow, N=12, 4 tomograms) lattice spacings. Scale bars: 100 nm **(A**,**D)**, 50 nm **(B**,**E)**. Untreated distribution is significantly different from Taxol distribution (p-value = 0.0006, unpaired t-test based permutation-test).

We first assessed the lattice spacing of microtubules from untreated U2OS cells using this *in situ* layer line analysis. In line with *in vitro* results of GDP-bound microtubules, 74% (23 out of 31) had a compacted average monomer spacing of 40.8-41.1 Å (Figure 1H). The remaining microtubules had an expanded spacing of 41.5-42.7 Å. These distinct lattice spacings could even be detected within the same tomogram (Supplemental Figure 2). Our results indicate that microtubule lattice spacings are diverse even far away from microtubule tips, where an altered lattice spacing is expected (Hyman et al., 1995; LaFrance et al., 2021). Within the field of view of our data, which is approximately 0.8 μm, we did not observe any microtubule ends. In contrast to the changes in lattice spacing at microtubule ends, which are important for the regulation of microtubule growth dynamics (Manka and Moores, 2018b), the different lattice spacings that we observe in the core of the microtubule likely play a role in microtubule stability, MAP binding and motor protein kinetics, as has been previously postulated (Cross, 2019).

**Figure 2.**
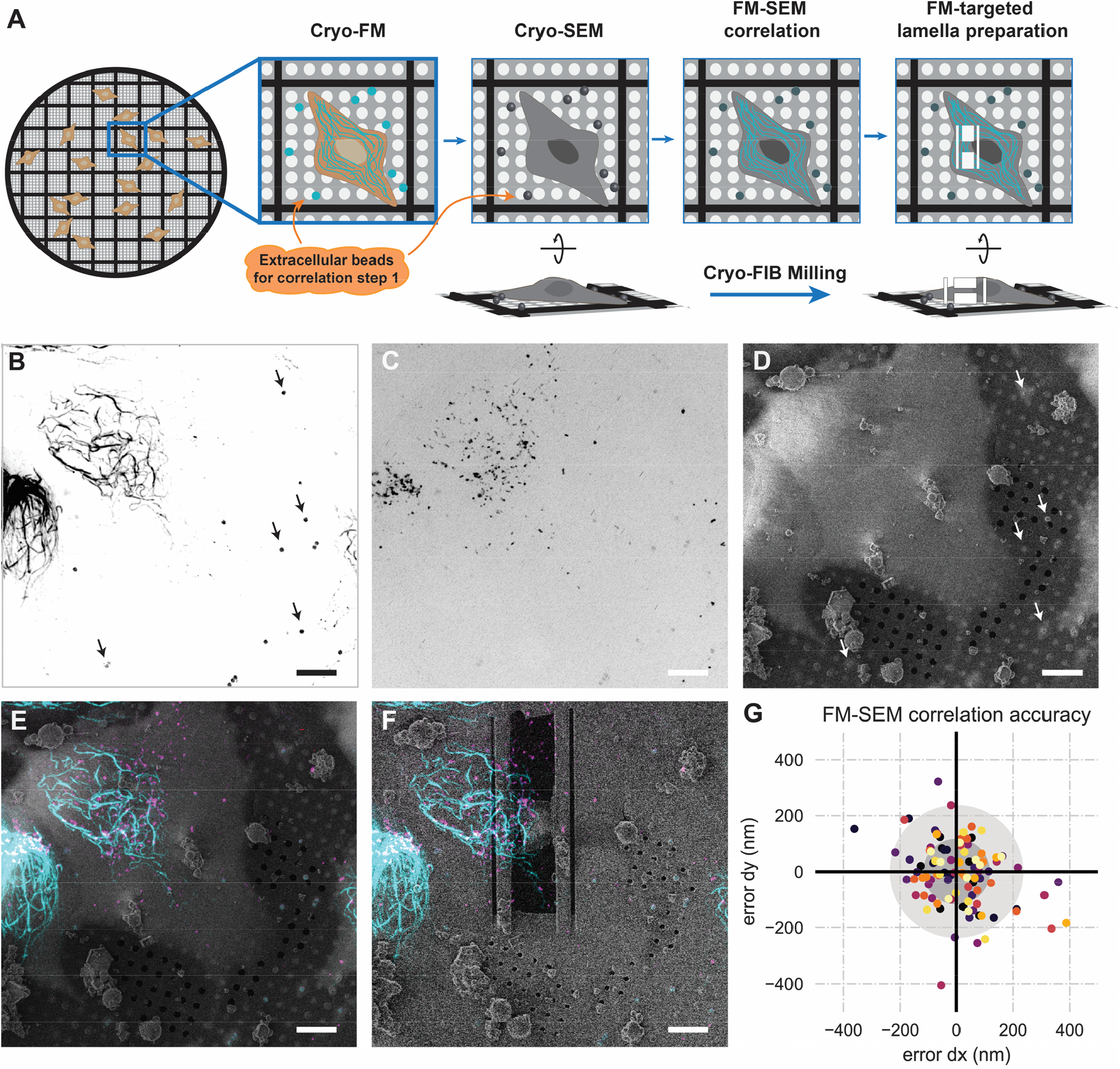
Correlation of FM to SEM data for targeted cryo-FIB milling. (**A**) Cartoon describing the FM-SEM correlation performed using extracellular beads. (**B**) MIP of a rigor-2xmNeongreen z-stack, beads used for 3D correlation are indicated with arrows. (**C**) MIP of a z-stack with fBSA-Au^5^ beads used for the subsequent FM-TEM correlation (see Figure 3). (**D**) Untilted SEM image of the same grid square as shown in B and C. (**E**) Correlated rigor-2xmNeongreen and fBSA-Au^5^ overlayed with the SEM image. (**F**) Correlated rigor-2xmNeongreen and fBSA-Au^5^ overlayed with the untilted SEM image of the polished lamella (milled at a 9 degree angle). (**G**) Scatterplot of correlation errors from leave-one-out calculations, each dataset has a unique colour, grey circles mark the 1xSD and 2xSD boundaries (12 datasets, 96 beads). Scale bars: 10 μm (**B-F**).

To confirm that our *in situ* layer line analysis is sensitive to changes in lattice spacing, we set out to modulate the microtubule lattice and test whether this change could be detected. Microtubules that are polymerized *in vitro* in the presence of Taxol are structurally altered compared to the GDP-bound, compacted state (Alushin et al., 2014; Kellogg et al., 2017; Rai et al., 2019). We therefore hypothesized that Taxol might expand microtubules inside cells as well. Indeed, in U2OS cells treated with Taxol we observed that most microtubules (92%, 11/12) have an expanded lattice spacing of 42.3- 43.2 Å (Figure 1D-H), significantly different from the untreated distribution (p-value = 0.0006). Together, these data show that changes in lattice spacing can be detected with our *in situ* layer line analysis and that the overall distribution of microtubule lattice spacings can be altered by Taxol.

Remarkably, the Taxol-induced expansion to 41.2-42.0 Å observed *in vitro* is considerably smaller than the hyper-expansion measured using our *in situ* approach (Alushin et al., 2014; Estevez-Gallego et al., 2020; Kellogg et al., 2017; Rai et al., 2019; Vale et al., 1994). A complex interplay between Taxol, MAPs and PTMs might exist as up or downregulation of MAPs, like Tau, can affect the sensitivity of cells to Taxol (Orr et al., 2003). Additionally, Taxol treatment leads to an increase in PTMs like acetylation (Hammond et al., 2010). These components are usually completely or partially absent from *in vitro* systems, which may explain why this hyper-expansion has thus far remained unreported.

### Fluorescence microscopy-guided cryo-FIB milling

Having established that different microtubule lattice spacings can be monitored within the cell, we next set out to determine the lattice spacing distribution of the subset of microtubules to which kinesin-1 selectively binds (Burute and Kapitein, 2019). To this end we used rigor-2xmNeonGreen, a fluorescently tagged mutant of kinesin-1 that has a very low rate of microtubule unbinding and binds to a subset of stable microtubules (Jansen et al., 2021). We set up a two-step cryo-CLEM workflow, wherein fluorescence microscopy (FM) data of U2OS Flp-In T-Rex cells expressing rigor- 2xmNeonGreen were used to target lamella preparation sites in the first correlation step (Figure 2A), and then used to distinguish the kinesin-1 bound microtubules from unbound microtubules in the lamella visible in the transmission electron microscope (TEM) in the second step (Figure 3A). For the first step, the extracellular beads visible in both the FM stack and the SEM image were used to overlay the FM stacks of the rigor-2xmNeonGreen (Figure 2B) and endocytosed fBSA-Au^5^ beads (Figure 2C) (Fermie et al., 2022) with the SEM image (Figure 2D). The resulting overlay (Figure 2E) was used for targeted milling (Figure 2F).

**Figure 3.**
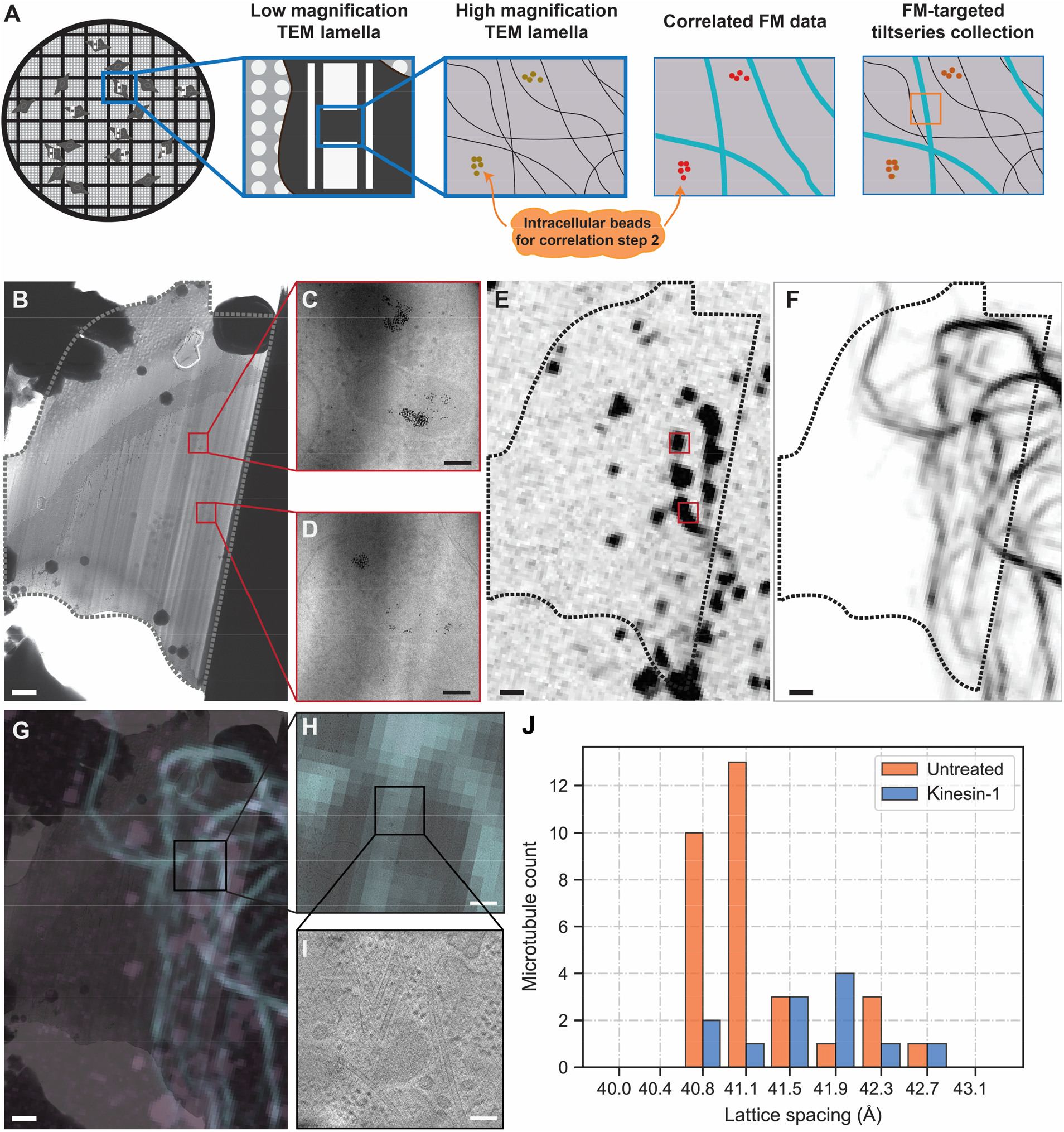
Kinesin-1 bound microtubules have a slightly expanded lattice compared to the GDP- compacted lattice. (**A**) Cartoon describing the FM-TEM correlation performed using intracellular beads. (**B**) TEM overview image of a lamella, dotted line indicates outline of the lamella, red squares the location of the fBSA-Au^5^ beads. (**C and D**) Zoom in of the fBSA-Au^5^ beads found both in the TEM lamella in B and in the correlated fBSA-Au^5^ FM data in E. (**E**) Correlated fBSA-Au^5^ FM data, red squares indicate fBSA-Au^5^ location. (**F**) Correlated rigor-bound microtubules. (**G**) Overlay of TEM lamella and correlated FM data of both fBSA-Au^5^ (pink) and rigor-bound microtubules (blue). (**H and I**) Two step zoom of two microtubules overlapping with the rigor-bound FM data. (**J**) Histogram showing the untreated lattice spacings (N=31, 12 tomograms, same data as Fig. 1H, included for comparison) and the lattice spacings of the kinesin-1 bound subset of microtubules (N=12, 6 tomograms). Scale bars: 1 μm (**B, E, F and G**), 100 nm (**C**,**D and I)**, 400 nm (**H**). Untreated distribution is significantly different from kinesin-1 distribution (p-value = 0.045, unpaired t-test based permutation test).

The FM-SEM localization accuracy was estimated using a “leave-one-out” metric (see materials and methods) and showed an FM-SEM correlation error with a standard deviation of 121 nm in x and of 116 nm in y (Figure 2G). The slight increase in accuracy that we obtained in comparison to the previous accuracy measurements obtained by Arnold et al. might be due to the increased number of beads we used for calculating the transform (Arnold et al., 2016). This estimated localization error confirmed that the FM-SEM correlation was sufficiently accurate for our CLEM approach.

### Kinesin-1-bound microtubules inside cells are expanded

Next, the correlated FM data were overlayed with the TEM overview image of the lamella (Figure 3A). To confirm that the FM-TEM correlation was successful, fBSA-Au^5^ beads present in the FM data were localized in the TEM image of the lamella (Figure 3B-E) and fine adjustments based on the higher resolution TEM data were performed. This was critical as the microtubules alone did not allow for assessment or optimization of the correlation (Figure 3F). When the correlation was considered unambiguous and reliable, the rigor-2xmNeonGreen data was used to localize the kinesin-1 bound microtubules within the lamella (Figure 3G-I). Analysis of their lattice spacings revealed a clear shift towards a more expanded lattice distribution, centred around 41.5-41.9 Å (p-value = 0.045), in comparison to the dominant compacted lattice observed for untargeted microtubules (Figure 3J). The observed distribution was also significantly different from the lattice distribution observed for Taxol treated cells (p-value = 0.0107). Our results indicate therefore that kinesin-1 bound microtubules inside cells are predominately more expanded than those not bound by kinesin-1.

The observed preference for an expanded lattice might relate to the structure of kinesin-1 or result from cooperative behaviour with other MAPs. The specific length of the neck linker of kinesin- 1 results in a notably high processivity compared to other kinesin motors (Shastry and Hancock, 2010; Yildiz et al., 2008). An expanded lattice might better accommodate the step size and increase processivity further. In these experiments the structural changes that occur upon microtubule expansion may result in an increased interaction and binding affinity between the L11-α4 junction of kinesin-1 and the H3 helix of α-tubulin (Morikawa et al., 2015; Shima et al., 2018). Alternatively, additional interactions with MAPs, such as MAP7 that may be sensitive to the lattice spacing, could ultimately be responsible for recruiting kinesin-1 (Ferro et al., 2022; Hooikaas et al., 2019; Monroy et al., 2020).

Despite the additional correlation accuracy gained with the intracellular beads, we cannot exclude that a proportion of the kinesin-1 bound microtubules were identified incorrectly due to the limited resolution of our cryo-FM setup. This fractional misassignment might contribute to the large spread observed in lattice spacings for the kinesin-1-bound microtubule-subset. Technical FM improvements such as cryo-MINFLUX (Gwosch et al., 2020), engineered point spread functions (Zhou et al., 2019) or integrated FM-FIB solutions (Bieber et al., 2021) may improve the cryo-FM resolution (lateral and/or axial) and resulting CLEM localization accuracy, as well as the throughput of future cryo-CLEM workflows. Nevertheless, the current workflow, which uses an additional independent marker (the fBSA-Au^5^ beads) for the FM-TEM correlation, can readily be used to answer a broad range of biological questions.

Taken together, our approach provides an unbiased way to investigate microtubule lattice spacings inside the cell. We found that most microtubules have a compacted lattice of around 41 Å, in line with previous *in vitro* and *in situ* studies (Figure 4, row I and II). Importantly, we observed that a range of expanded lattices can be found within the cell. Thus far, expanded states have only been reported in *in vitro* studies when microtubules are assembled using the GTP analogue GMPCPP or the drug Taxol (Figure 4, row III and IV). The relevance of different lattice spacings for MAP behaviour has only been investigated for a few examples and expanded lattices have mostly been linked to plus-end regulating proteins (Figure 4, row V). However, our data, together with *in vitro* data, indicate that kinesin-1 preferentially binds to an expanded lattice (Peet et al., 2018; Shima et al., 2018). The microtubule lattice might therefore function as an additional mechanism to guide protein binding specificity, on top of the previously established elements of the tubulin code. Similar analysis on other members of the kinesin family may reveal the extent to which these motors are sensitive to the lattice spacing of microtubules.

**Figure 4.**
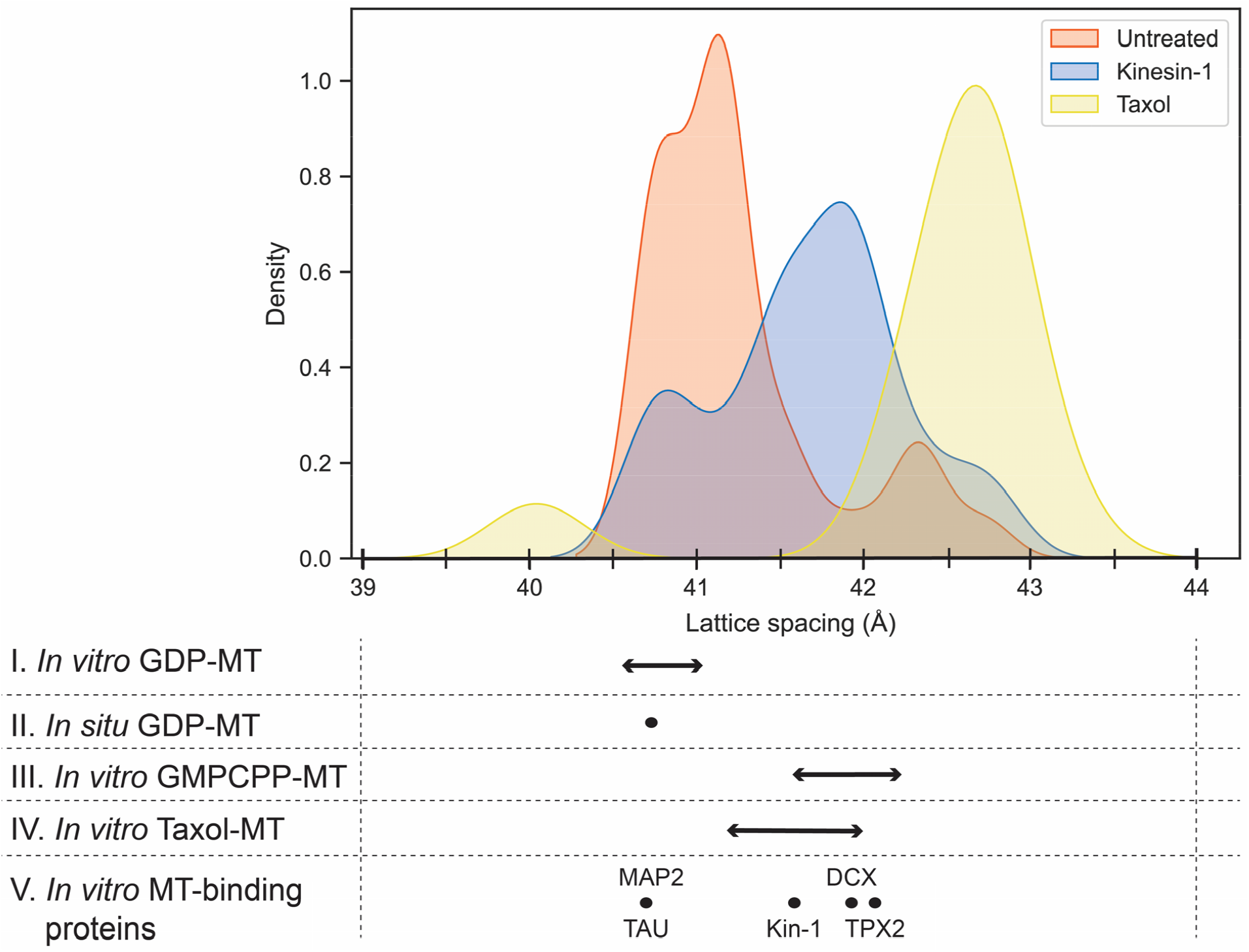
*In situ* microtubule landscape in relation to previously reported lattice spacings. Upper panel shows a density distribution calculated from the different lattice spacing datasets measured here. The table below shows the reported lattice spacings for the GDP lattice both *in vitro* and *in situ* (row I and II), and for *in vitro* GMPCPP and Taxol-bound lattices (row III and IV). Row V shows MT-binding proteins for which binding in relationship to the microtubule lattice spacing has been investigated. References can be found in supplementary table 1.

In this paper we present a two-step cryo-CLEM workflow, which we used to study microtubule lattice spacing within unperturbed cells, upon Taxol treatment, and in correlation with kinesin-1 binding. The discovery of diverse microtubule lattice spacings within the cell emphasizes the need to further investigate the relationship between the microtubule lattice and protein binding behaviour. Furthermore, this study shows the potential of our cryo-CLEM workflow to gain new insights into well-studied biological processes from intact cells.

## Acknowledgements

We thank Alessandro Sartori (Institute of Molecular Cancer Research, Switzerland) for providing us with the U2OS Flp-In T-Rex stable cells, Vladan Lucic and Mihajlo Vanevic for computational support, Mariska Gröllers-Mulderij for guidance in cell culture, Rutger Hermsen for the input on statistical data analysis, Mathieu Baltussen for help with figure design and Nalan Liv for providing us with the intracellular fBSA-Au^5^ beads. This work was supported by the ERC Consolidator grant 724425 (Biogenesis and Degradation of Endoplasmic Reticulum Proteins, to F.F.) and ERC Consolidator grant 819219 (to L.C. Kapitein), and was part of the research programme National Roadmap for Large-Scale Research Infrastructure 2017 – 2018 with project number 184.034.014, which is (partly) financed by the Dutch Research Council (NWO). K.I. Jansen was funded by NWO (NWO-Graduate program project 022.006.001 to K.I. Jansen). The Netherlands Electron Microscopy Infrastructure (NEMI) helped support access to the Netherlands Center for Electron Nanoscopy (NeCEN) with support from operators Wen Yang and Willem Noteborn.

## Materials and Methods

### List of reagents

List of reagents:
| Reagent or resource | Source | Identifier |
| --- | --- | --- |
| **Q5 HF DNA polymerase** | NEB | Cat# M0491L |
| **Gibson Assembly® Master Mix** | NEB | Cat# E2611L |
| **DMEM + GlutaMAX-I** | GIBCO | Cat# 61965-026 |
| **DPBS** | Corning | Cat# 21-031-CV |
| **Pen Strep** | GIBCO | Cat# 15140-122 |
| **Trypsin-EDTA** | Gibco | Cat# 25200-056 |
| **Fetal Bovine Serum** | Sigma-Aldrich, Inc. | Cat# F7525-500ML |
| **Fibronectin** | Sigma-Aldrich, Inc. | Cat# F1141-2MG |
| **Blasticidin** | Thermo Fisher Scientific | Cat# R21001 |
| **Hygromycin B** | Corning | Cat# 30-240CR |
| **Paclitaxel/Taxol** | Thermo Fisher Scientific | Cat# P3456 |
| **Doxycycline** | Sigma-Aldrich, Inc. | Cat# 24390-14-5 |
| **fBSA-Au<sup>5</sup>** | Cell Microscopy Core, Utrecht UMC | N.A. |
| **Dynabeads™ MyOne™ Carboxylic Acid** | Thermo Fisher Scientific | Cat# 65011 |
| **Cellview cell culture dish** | Greiner bio-one | Cat# 627860 |
| **Quantifoil 200 mesh holey carbon R2/2 gold grids** | Quantifoil Micro Tools | N.A. |
| **Whatman Filter Paper** | Whatman | Cat# 10311610 |

### Cell lines and cell culture

U2OS wild type (WT) cells were purchased from ATCC and U2OS Flp-In T-Rex cells were a kind gift from Prof. Alessandro Sartori (Institute of Molecular Cancer Research, University of Zurich). Cells were confirmed to be free of mycoplasma. The U2OS Flp-In cell line that upon doxycycline-induction expresses hKif5b(1-560)G234A-mNeongreen-mNeongreen (hereafter referred to as rigor- 2xmNeongreen) (Jansen et al., 2021) was derived from the U2OS Flp-In cell line by transfection with the pCDN5/FRT/TO vector (Invitrogen) and pOG44 vector (Invitrogen). The U2OS rigor- 2xmNeongreen cell line was cultured in Dulbecco’s Modified Eagle’s Medium + GlutaMAX^™^-l (DMEM-Glu) supplemented with 10% FBS, 100 U/ml penicillin, 100 μg/ml streptomycin, 15 μg/ml blasticidin S and 0.25 mg/ml hygromycin B. To induce expression of rigor-2xmNeongreen, doxycycline (10 ng/mL) was added to the cells 24 hrs before plunging. U2OS WT cells were cultured in DMEM- Glu supplement with 10% FBS, 100 U/ml penicillin and 100 μg/ml streptomycin. Cells were kept at 37°C and 5% CO_2_.

### Plasmids and cloning

To generate a stable, isogenic U2OS Flp-In cell line, rigor-2xmNeongreen was subcloned into pCDNA5/FRT/TO (Invitrogen) via Gibson Assembly using the primer set ‘5-GCTCGGATCCACTAGTCCAGTGTGGTGGAATTCTGCAGATGCCACCATGGCGGACCT-3’ and 5’-ACGGGCCCTCT AGACTCGAGCGGCCGCCACTGTGCTGGATGCGGCCGCTTACTTGTACAG-3’. The G234A rigor mutation was used as initially described (Rice et al., 1999). mNeongreen (Shaner et al., 2013) was provided by Allele Biotechnology. The FLP recombinase expression vector is encoded in pOG44 (Invitrogen).

### Sample preparation

Quantifoil 200 mesh holey carbon R2/2 gold grids (Quantifoil Micro Tools) were glow discharged (PELCO easiGlow, Ted Pella) and placed on 40 μL fibronectin droplets (50 μg/μL) and incubated at 37°C for 2-3 hrs. Subsequently, the grids were washed two times by placing them on droplets of 40 μL PBS and put in 35 mm glass bottom dishes (Greiner Bio-One). 90.000 cells in 2 mL medium were seeded on these grids and were left to settle in the hood for ∼20 min. before placing them in the incubator. After 24 hrs, the U2OS rigor-2xmNeongreen cells were treated with doxycycline (10 ng/mL) while WT U2OS cells were treated with Taxol (1 μM), if applicable. Plunging was performed 48 hrs after cell seeding. In preparation for vitrification, the cells were incubated with fBSA-Au^5^ (diluted to OD 5) for 3-4 hrs (Fermie et al., 2022). Just before plunging, the media was exchanged for fresh DMEM. The grids were washed by dipping 2 times in PBS (37°C) before 3 μL of 1 μm Dynabeads (Thermo Fisher Scientific: MyOne with 40% iron oxide, carboxylic acid) diluted 1:20 in PBS, was added to the grids. Finally, the cells were vitrified in liquid ethane after manually blotting for 10s. The grids were clipped into autogrids and kept at liquid nitrogen temperature throughout the subsequent experiments.

### SEM grid screening

To increase efficiency, although at the cost of ice contamination, grids were screened in the cryo-FIB SEM (Aquilos^™^, Thermo Fisher Scientific), prior to fluorescent imaging. SEM grid overview images were taken using FEI MAPS 3.8 software. Grids with clearly visible grid holes and an appropriate distribution of cells were used in the next steps of the cryo-CLEM workflow.

### Cryo-fluorescence microscopy

Cryo-FM data were obtained with the FEI CorrSight™ equipped with a cryo-stage, using the Andromeda spinning-disk confocal microscope module. Grid overviews were collected with a 5x/0.16 NA air objective using transmitted light. SEM overview images were aligned to the FM overview images via 3-point correlation in the MAPs v3.8 software (Thermo Fisher Scientific). Based on the SEM and FM overlay, candidate cells were selected. Using the EC “Plan-Neofluar” 40x/0.9 NA air objective, z-stacks ranging from 8-14 μm with 300 nm step size were collected. Z-stacks were recorded to capture 3 different fluorescent probes, namely rigor-2xmNeongreen (488 nm), Dynabeads (488 nm) and fBSA-Au^5^ (561 nm). Images were recorded with FEI MAPS v3.8 software and LA FEI Live Acquisition v2.2.0 (Thermo Fisher Scientific). The images were subjected to a mild deconvolution using Huygens Professional software v 21.04 (Scientific Volume Imaging) with classic maximum likelihood estimation algorithm.

### Targeted cryo-focussed ion beam milling

Cryo-FIB milling was performed in the cryo-FIB SEM (Aquilos^™^, Thermo Fisher Scientific). Lamellae were prepared as described previously (Wagner et al., 2020). SEM grid overview images were obtained and overlayed with the cryo-FM overview images using 3-point correlation available in the MAPS v3.8 software to guide the localization of candidate lamella sites. The grids were coated with platinum to reduce charging effects. Eucentric heights and minimal stage tilt angles (16-18°, corresponding to 9- 11° lamella angle), to ensure access of the ions to the milling sites, were determined. High magnification SEM images (0.135 μm/pixel, 1 μs dwell time, 1,536×1,024 pixels, 2 keV, 13 pA) of each target cell were taken and manually overlayed with the corresponding cryo-FM maximum image projection (MIP) using Dynabeads as fiducials. The location of the milling patterns was based on the correlated cryo-FM data. An FM-FIB correlation was not included in our workflow as this is a time-intensive procedure - and the limited z-resolution of our cryo-FM set up meant this correlation did not add significant information when tested.

Next, the grids were subjected to organo-platinum deposition for 10 s via an integrated gas injection system to generate a more even surface and thereby reduce curtaining effects and protect the final lamella. Milling was performed with a stepwise decreasing current of 1 to 0.3 to 0.1 nA, and a shrinking milling pattern. The final polishing step was performed at 30 pA to reach a final lamella thickness of 80-140 nm. High magnification SEM images of the polished lamellae were taken (0.135 μm/pixel, 300 ns dwell time, 1,536×1,024 pixels, 2 keV, 13 pA) and the grids were coated with platinum for a second time before unloading.

### Targeted cryo-electron tomography

In preparation for FM-guided cryo-electron tomography data collection, SEM images of polished lamellae were aligned to the corresponding cryo-FM z-stacks, guided by high-dose SEM images of the lamella sites. Using the 3D Correlation toolbox (Arnold et al., 2016) 6-9 Dynabeads were localized both in the SEM image and the FM z-stack and a transformation matrix was fitted for the two sets of X,Y,Z coordinates, while aiming for an RMSD smaller than 1 pixel for each bead. Z-stacks were transformed according to this matrix with the 3D rigid body transformation of the Pyto python package (Arnold et al., 2016). MIPs of the transformed z-stack of each channel were overlayed with the SEM image of the polished lamella in FIJI (Schindelin et al., 2012) to generate a correlated FM-SEM overlay.

Lamellae were imaged on a 200 kV Talos Arctica transmission electron microscope (TEM) (Thermo Fisher Scientific) with a K2 summit direct electron detector (Gatan) or on a 300 kV FEI Titan Krios TEM (Thermo Fisher Scientific) with a K3 summit direct electron detector (Gatan), both equipped with a post-column energy filter aligned to the zero-loss peak and a 20 keV slit width. Using SerialEM (Mastronarde, 2003), lamellae overview images were taken at 7,300x (Arctica, 18.7 Å/pixel) or 4,800x (Krios, 38.6 Å/pixel).

The TEM overview image of the lamella and the corresponding FM-SEM overlay were further manually correlated. This last correlation step was guided by the bimodal intracellular fBSA-Au^5^ fiducials. In cases where the rigor-2xmNeongreen FM signal overlapped with a microtubule in the cryo-TEM lamella image, a tilt series was collected. The microtubules in the lamellae of untreated and Taxol treated cells were chosen at random. Tilt series of microtubules were recorded at a pixel size of 2.17 Å/px, a dose rate of ∼5 (Arctica) or 10-20 (Krios) e^−^/pixel/s and a total dose of 90-100 e-/Å^2^. All tilt series were collected using a dose symmetric scheme (Hagen et al., 2017), a tilt increment of 2° or 3°, a defocus target of -2.3 μm, and a tilt range of 69° to -51° or 51° to -69°, depending on the lamella orientation in the microscope.

### Tomogram reconstruction

The tilt series were aligned and dose-weighted using MotionCor2 (Zheng et al., 2017). The tilt series were generally aligned via patch tracking. If fBSA-Au^5^ beads were present, fiducial tracking was used. Tomogram reconstruction was performed in Etomo, part of the IMOD 4.10.29 package (Kremer et al., 1996). Contrast transfer function estimation and correction was performed in IMOD using the *ctfplotter* and *ctfphaseflip* commands and the tomograms were reconstructed using weighted back-projection and a SIRT-like filter with 3 iterations.

### Layer line analysis

Reconstructed tomograms were loaded in Dynamo (Castano-Diez et al., 2017). The filament model (crop along axis) was used to pick the microtubule backbone. The backbone coordinates were exported from MATLAB and used to generate a soft mask around the microtubule [in house script, available on GitHub at ldejager/InSitu_LayerLine_Analysis]. Particles with a box size of 190 pixels were extracted from the unbinned, masked tomograms. Microtubule particles were aligned in Dynamo using a reference based on a previously deposited microtubule structure (EMD-10896). After alignment, the particles were inspected using the *dGallery* command. Per microtubule, multiple aligned particles were selected to cover the whole microtubule. These particles were extracted from the unbinned tomograms with a box size of 1,030 pixels and summed along the z-axis [in-house script, available on GitHub at ldejager/InSitu_LayerLine_Analysis]. The power spectrum of each particle was calculated in FIJI. The power spectra of the particles from the same microtubule were summed to increase the signal-to-noise ratio. Layer lines were localized by calculating a line profile plot of the power spectrum. The average lattice spacing of the microtubule was calculated using formula 1.

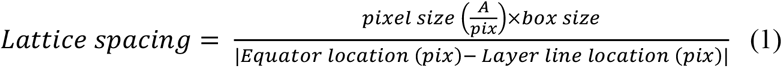

### Statistical analysis

Leave-one-out calculations were performed as described by (Schorb and Briggs, 2014). Briefly, a transformation matrix was calculated with a set of beads. This transform was applied to a bead not included in the initial set. The deviation of the predicted bead position from the true bead position was used as an accuracy measurement. P-values for comparison of the different lattice spacing distributions were calculated with a unpaired two-tailed t-test based permutation test with 10,000 iterations. Lattice spacing density distribution (figure 4) was calculated as a gaussian kernel density estimate. Permutation tests were performed in and graphs (histograms and distribution) were created with Jupyter notebook 6.0.3 (Kluyver et al., 2016).

**(Standard abbreviations: GTP, GDP, NA, WT)**

## SUPPLEMENTARY MATERIALS

**Supplementary Figure 1.**
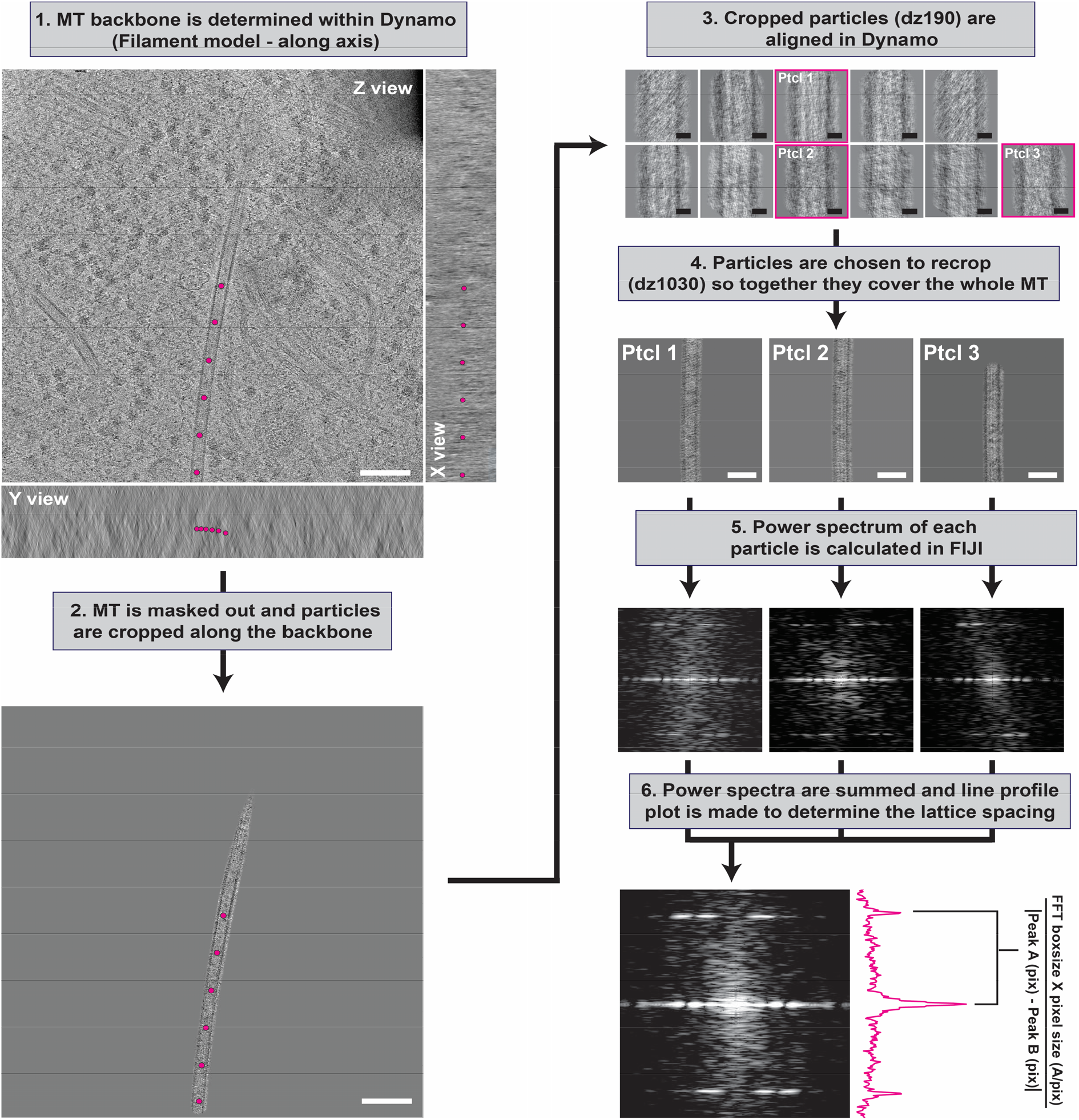
*In situ* layer line analysis workflow. Graphical depiction of the steps performed during the layer line analysis. (**Step 1**) Microtubule backbone is picked in dynamo. (**Step 2**) Using the backbone coordinates, a mask around the microtubule is generated and particles are cropped. (**Step 3**) The cropped particles (box size 190) are aligned and picked so that after recropping (**step 4**) with a box size of 1030 the whole microtubule is covered. (**Step 5**) 2D power spectrum of the MIP of each particle is calculated. (**Step 6**) power spectra of all particles part of the same microtubule are summed and the final power spectrum is used to localize the layer line and thereby calculate the lattice spacing. Scale bars: 100 nm (step 1, 2), 10 nm (step 3), 50 nm (step 4).

**Supplementary Figure 2.**
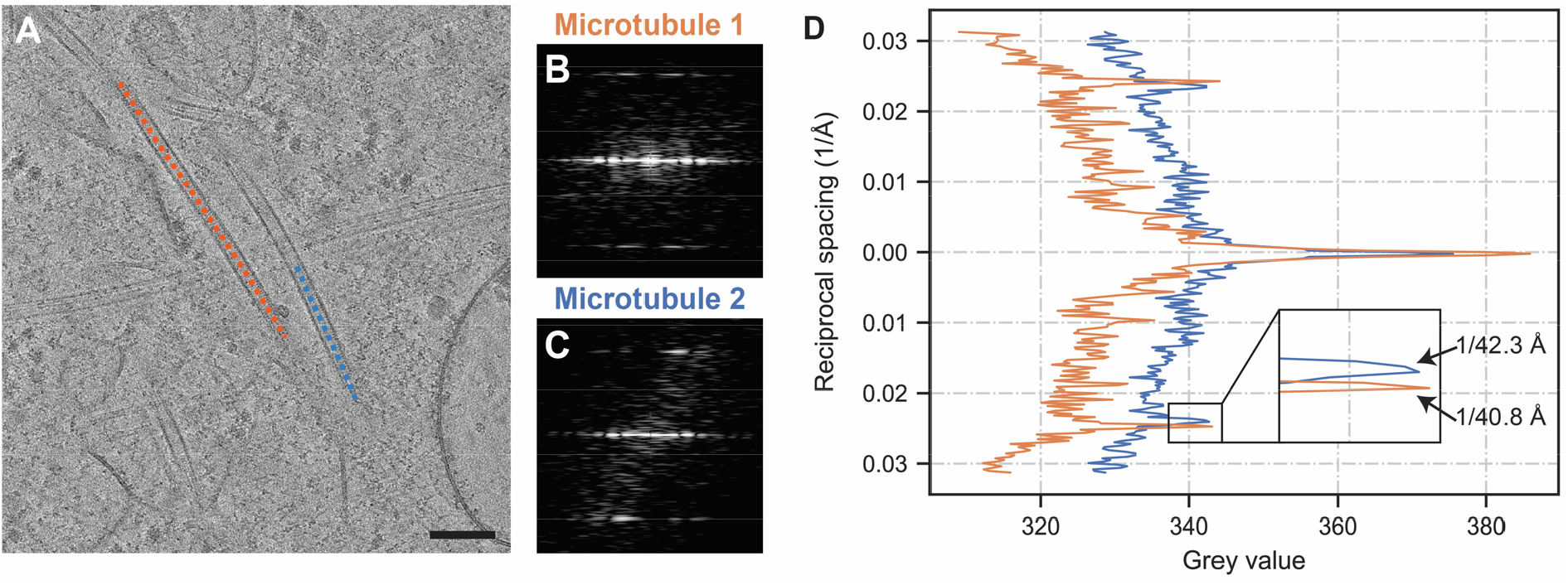
Expanded and compacted microtubule lattices within one tomogram. Tomogram where two MT backbones were analysed in parallel. (**A**) MT backbones were localized and processed as explained in Supplemental Figure 1. (**B and C**) show the final, summed power spectra of MT1 (orange) and MT2 (blue). (**D**) Layer line plot of the summed power spectra of MT1 and MT2. Scale bar: 100 nm (**A**).

**Supplementary Figure 3.**
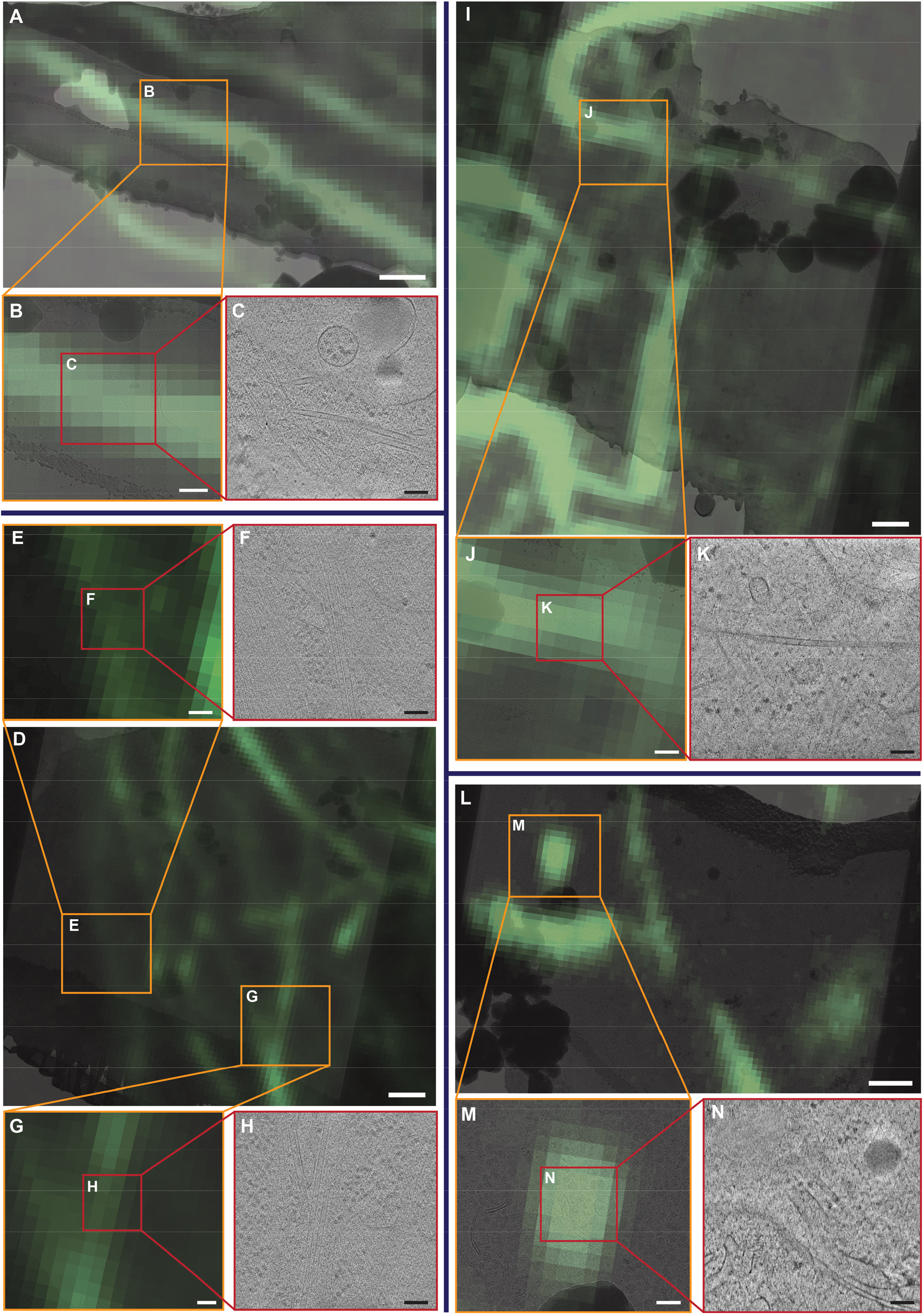
Kinesin-1 bound microtubules found via our cryo-CLEM set up. (**A**) TEM lamella overlaid with correlated rigor-m2xNeonGreen cryo-FM data. (**B and C**) One time and two time zoom in of the region of interest with the correlated microtubules. (**D-F, D/G/H, I-K and L-N**) Similar examples of correlated microtubules. Scale bars: 1 μm (**A, D, I, L**), 300 nm (**B, E, G, J, M**) and 100 nm (**C, F, H, K, N**).

**Supplementary table 1.**
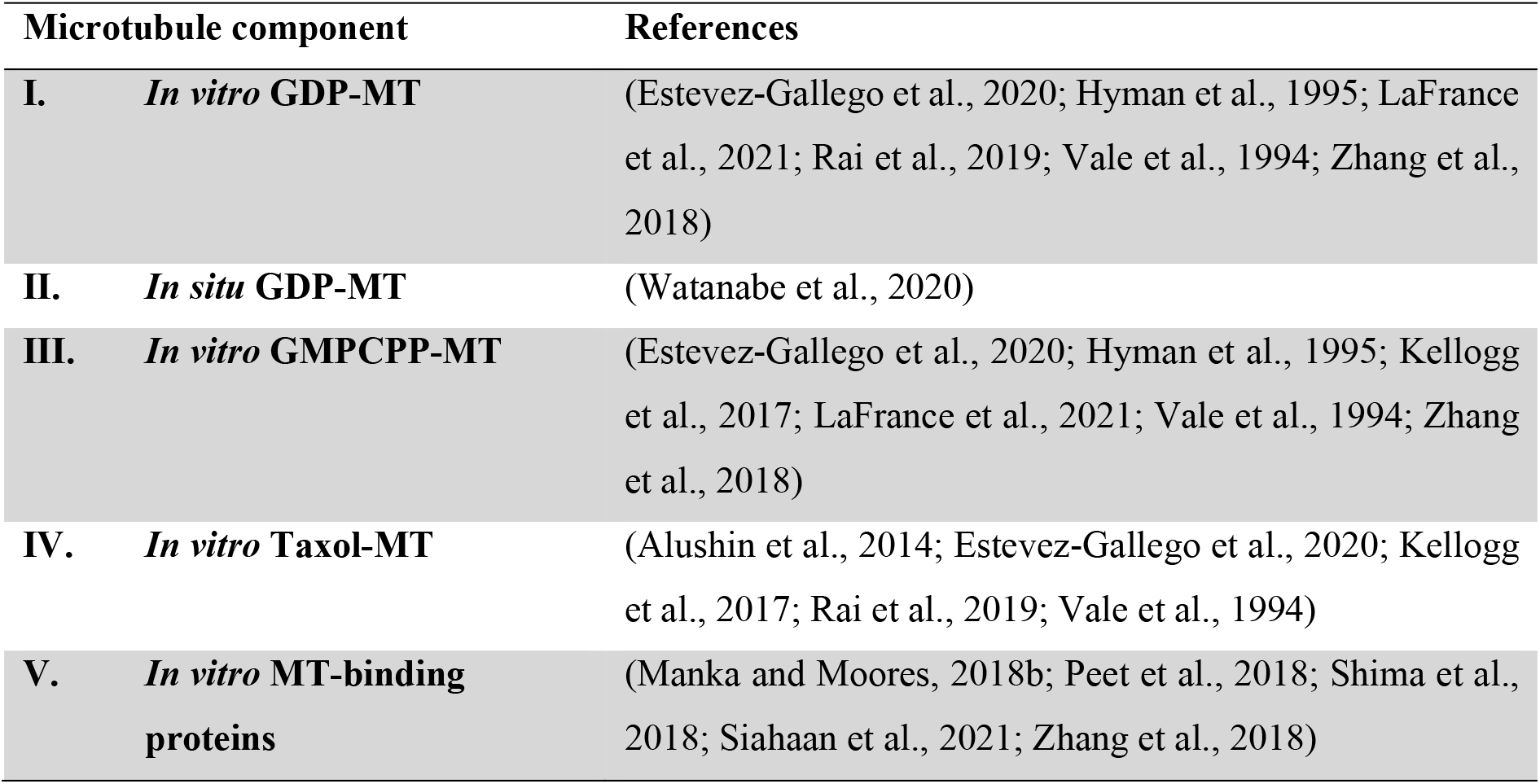
Literature that investigates the microtubule lattice spacings with respect to nucleotide state, Taxol treatment and MT-binding proteins.

## List with abbreviations

**Abbreviation Meaning**

CLEM: Correlative light and electron microscopy
GMPCPP: Guanylyl-(α,β)-methylene-diphosphonate
FIB: Focused ion beam
FM: Fluorescence microscopy
MAP: Microtubule associated proteins
MIP: Maximum image projection
MT: Microtubule
PTM: Post-translational modification
SEM: Scanning electron microscopy
TEM: Transmission electron microscopy

**(Standard abbreviations: GTP, GDP, NA, WT)**

